# Diffusion and interaction dynamics of the cytosolic peroxisomal import receptor PEX5

**DOI:** 10.1101/2021.05.25.445571

**Authors:** S Galiani, K Reglinski, P Carravilla, A Barbotin, I Urbančič, J Ott, J Sehr, E Sezgin, F Schneider, D Waithe, P Hublitz, W Schliebs, R Erdmann, C Eggeling

## Abstract

Measuring diffusion dynamics in living cells is essential for the understanding of molecular interactions. While various techniques have been used to explore such characteristics in the plasma membrane, this is less developed for measurements inside the cytosol. An example of cytosolic action is the import of proteins into peroxisomes, via the peroxisomal import receptor PEX5. Here, we combined advanced microscopy and spectroscopy techniques such as fluorescence correlation spectroscopy (FCS) and super-resolution STED microscopy to present a detailed characterization of the diffusion and interaction dynamics of PEX5. Among other features, we disclose a slow diffusion of PEX5, independent of aggregation or target binding, but associated with cytosolic interaction partners via its N-terminal domain. This sheds new light on the functionality of the receptor in the cytosol. Besides specific insights, our study highlights the potential of using complementary microscopy tools to decipher molecular interactions in the cytosol via studying their diffusion dynamics.

**Summary:** The peroxisomal import receptor PEX5 transports newly synthesized proteins from the cytosol to the peroxisomal matrix. Here the cytosolic diffusion and interaction dynamics of PEX5 are characterized by advanced microscopic spectroscopy methods, revealing a so far unknown interaction partner.

## Introduction

Cellular signaling critically depends on accurate interaction between molecules, and alterations may lead to severe cellular dysfunctions. Therefore, it is essential to employ observation techniques that disclose details of interaction dynamics in the living cells with high accuracy. One remedy is to study molecular diffusion dynamics, since diffusion will be hampered upon interactions (Blouin et al., 2016). Various fluorescence microscopy approaches have been employed and optimized to detail molecular diffusion dynamics especially in the cellular plasma membrane, such as Fluorescence Recovery After Photobleaching (FRAP) (Ladha et al., 1996), fluorescence correlation spectroscopy (FCS) (Schwille et al., 1999) or single-particle tracking (SPT) (Kusumi et al., 2005; Schütz et al., 1997), even in combination with super-resolution microscopy approaches (Eggeling et al., 2009a; Manley et al., 2008). However, employing these approaches to processes in the cytosol of the living cell is much more challenging, especially due to adding one spatial dimension from two-(2D) to three-dimensional (3D) diffusion. Therefore, the application of the above techniques to cytosolic studies is much more elaborate, usually requiring much data mining (Fritzsche and Charras, 2015; Kues et al., 2001; Wachsmuth et al., 2000; Wachsmuth et al., 2003). Therefore, it is necessary to further adapt techniques for studying cytosolic interaction dynamics, but also to combine them with dedicated complementary tools and controls.

As an example, organelle functions naturally rely on molecular interactions in the cellular cytosol, such as the import of proteins into peroxisomes. Peroxisomes are ubiquitous organelles in eukaryotic cells fulfilling many metabolic functions that are cell-type specific and variable as a response to environmental changes. Consequently, the pool of peroxisomal matrix proteins needs to be continuously adapted, entailing the necessity of a highly dynamic import system. Peroxisomal matrix proteins are synthesized on free ribosomes in the cytosol and transported into the organelle post-translationally. The peroxisomal cargo receptor PEX5 is one of the key proteins in the peroxisomal import process (Figure 1). Most peroxisomal matrix proteins imported by PEX5 contain a peroxisomal targeting signal type 1 (PTS1) at their C-terminus, while cargo proteins with the less abundant PTS2 targeting sequence are recognized and transported by the PTS2-receptor PEX7. PEX5 appears as two splice variants, a shorter one (PEX5S) that can only recognize PTS1 cargo proteins, and a longer variant (PEX5L) that contains an additional PEX7 binding site (Schliebs and Kunau, 2006). Therefore, the import pathways of PTS1 and PTS2 proteins merge with the long splice form. After binding, PEX5 directs the cargo-receptor complexes to the peroxisomal membrane and initiates the cargo translocation by interacting with the peroxisomal membrane protein PEX14. At the peroxisomal membrane, PEX5 gets integrated into the membrane, forming a transient translocation pore to import the cargo protein into the peroxisome (Meinecke et al., 2010). Consequently, PEX5 is a shuttling receptor with a much larger fraction in the cytosol (searching to bind newly synthetized cargo proteins) and only a small fraction at a time bound to the peroxisomal membrane (mainly involved in cargo-translocation) (Dodt and Gould, 1996; Galiani et al., 2016). While it has been shown that the import of cargo proteins depends on the affinity of the PTS1 signal sequence to PEX5 (Ghosh and Berg, 2010; Reglinski et al., 2015), no further details are known on the interaction time scales and thus dynamics, i.e. the diffusion dynamics of the cargo receptor complex in the cytosol are basically completely unknown, which yet can highlight important details of the involved interaction dynamics.

**Figure1:**
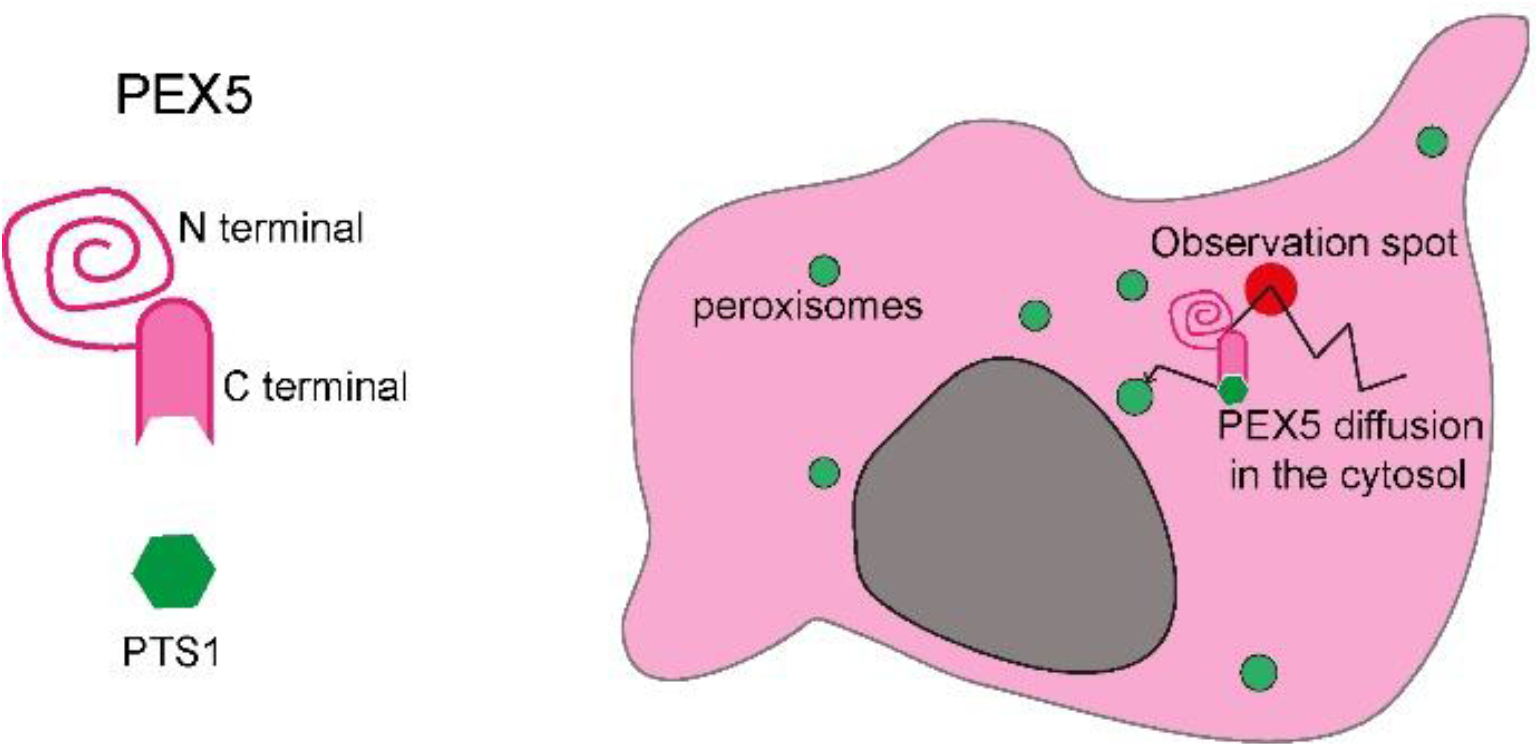
PEX5 receptor. **Left:** Schematic of the PTS1 receptor PEX5 and its binding partner, a PTS1 cargo-protein. PEX5 consists of a globular C-terminal domain, which binds the PTS1-containing peroxisomal matrix proteins and an unstructured N-terminal domain which is needed for peroxisomal docking, integration of PEX5 into the peroxisomal membrane, and translocation of the PTS1-cargo protein across the membrane. **Right:** Schematic of a cell expressing full length PEX5 (magenta) and eGFP-PTS1 (green at the peroxisomes). The diffusion (represented via the black arrow) of PEX5 and the PEX5/eGFP-PTS1 receptor/cargo complex is studied in the cytosol. Here the fluorescence fluctuations are measured in an observation spot (red).

Here, we present a detailed characterization of the diffusion and thus interaction dynamics of human cytosolic PEX5 *in vitro* and in living cells by combining state-of-the-art microscopy and spectroscopy techniques such as FCS in combination with multi-color detection and super-resolution microscopy together with biological manipulation such as CRISPR/Cas9 and model systems. As a result, we prove a free diffusion for PEX5 in the cytosol, which was found to be unexpectedly slow and independent of cargo binding. Among many controls, we investigate PEX5 oligomerization, interactions with other proteins of the PTS2 import pathway, binding to constituents of the peroxisomal membrane or association with the cytoskeleton, and we could show that none of this shows an influence on PEX5 diffusion. Interestingly, the slow diffusion of PEX5, which depends on its N-terminal half, is not linked to the intrinsically disordered structure of this region. By using the cell-derived giant plasma-membrane vesicle (GPMV) model system and recombinant proteins in solution, we show that the slow diffusion of PEX5 only occurred in the presence of cytosolic components, indicating that the characteristic diffusion of PEX5 is strictly linked to yet to be identified cytosolic factors.

## Results

### Wild type (PEX5L-SNAP) but not PTS1 non-binding mutant (PEX5L S600W-SNAP) shows co-diffusion with PTS1

To investigate the mobility of human PTS1 receptors and peroxisomal proteins in the cytosol, we expressed them in fusion with fluorescent proteins or with a SNAP-tag, which allows covalent binding of a fluorescent dye, in human fibroblasts (GM5756-T), and measured their diffusion and interaction characteristics by fluorescence correlation spectroscopy (FCS) and two-color fluorescence cross-correlation spectroscopy (FCCS) on a confocal microscope. FCS provides information on the molecular diffusion by recording the fluorescence signal over time and analyzing the fluctuations caused by stochastic movement of fluorescent molecules in and out of the confocal observation volume. From the autocorrelation function (ACF) of these fluctuations, it is possible to extract the average transit time of a protein of interest through the observation volume, and to determine its characteristic diffusion coefficient D and thus mobility as well as changes due to potential molecular interaction dynamics. In FCCS, the cross-correlation function (CCF) of the characteristic fluctuations of two differently labelled proteins is determined, highlighting their co-diffusion in addition to the mobility (Figure 2a).

**Figure 2:**
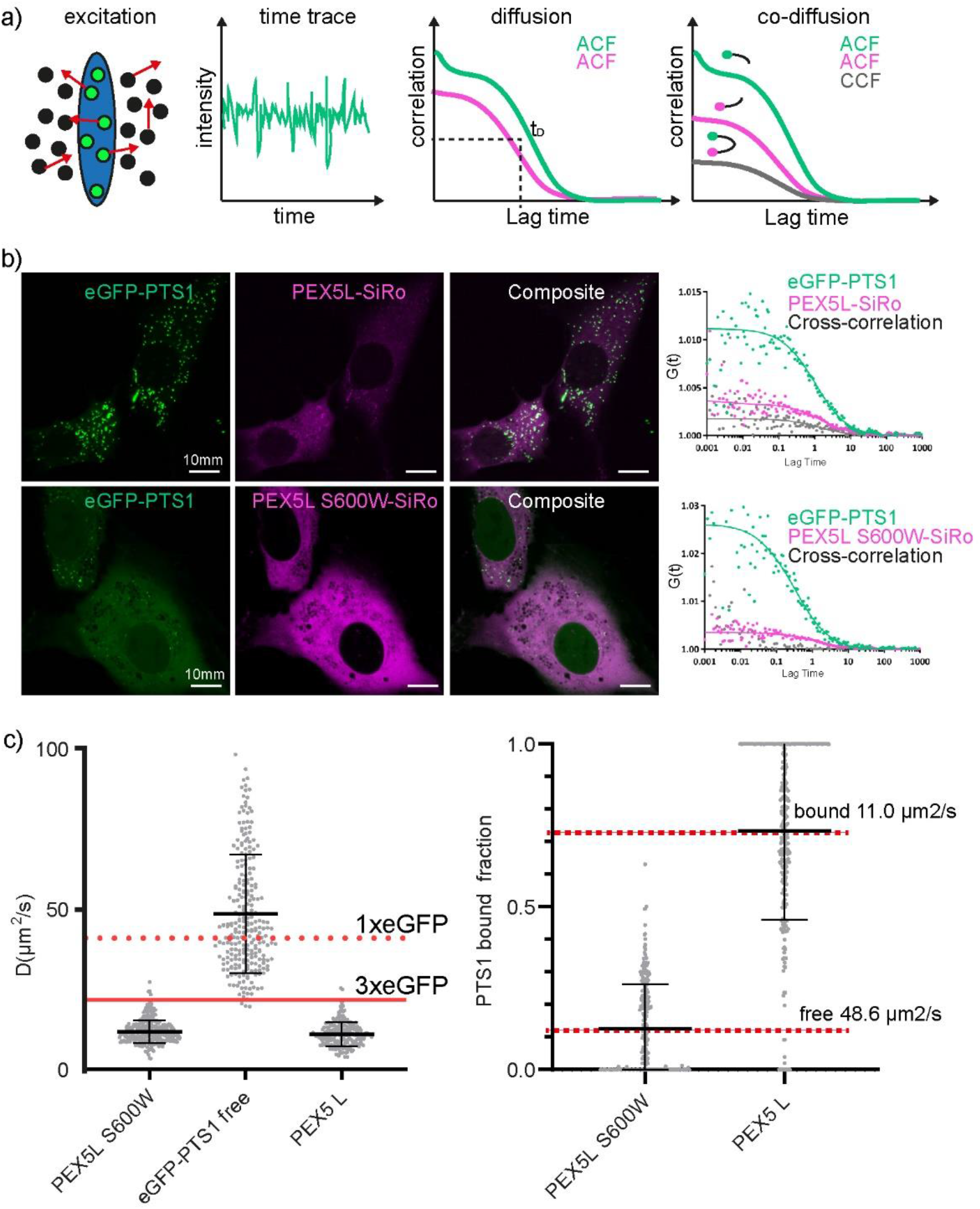
Characterization of the diffusion of PEX5L and eGFP-PTS1 by FCS. a): Principle of FCS and FCCS. FCS provides the average transit time, T_D_, of diffusing fluorescent molecules (green) through the microscope’s observation volume (blue) via calculation of the ACF from the time trace of the fluorescence signal. FCCS measures the interaction between two differently species of molecules diffusing through the observation volume by calculation of the cross-correlation function (CCF) between the two time traces. b): Representative images of human fibroblasts expressing eGFP-PTS1 (green) and PEX5L-SNAP-SiRo (magenta) (upper panel) and PEX5L S600W (lower panel) 24 h after transfection. The graphs on the right show the correlation curves (G(t), (dots), and fits (solid lines), of each individual species (eGFP-PTS1, PEX5L and PEX5L S600W) and their cross-correlation. Only PEX5L and eGFP-PTS1 signals cross-correlate (grey curve). c): Left: Diffusion coefficients (D) calculated from the fit of autocorrelation curves. Only one diffusing population could be calculated for free PEX5L S600W (11.9 ± 3.6 μm^2^/s), as well as mainly a free eGFP-PTS1 component could be calculated (48.6 ± 18.6 μm^2^/s). In cells expressing PEX5L only one slow diffusing population was found for PEX5L (11.0 ± 3.7 μm^2^/s), diffusing at the same speed as PEX5L S600W, independent from PTS1-cargo proteins interaction. Right: Fractions of eGFP-PTS1 diffusing together with PEX5L. eGFP-PTS1 was co-expressed with PEX5L S600W or PEX5L. In the case of PEX5L S600W most of the eGFP-PTS1 diffused freely and only a 12% of eGFP-PTS1 is bound to the endogenous PEX5. When PEX5L was expressed together with eGFP-PTS1, two diffusing populations (one bound to PEX5L and another free) for eGFP-PTS1 were found (11.0 ± 3.7 μm^2^/s and 48.6 ± 18.6 μm^2^/s, respectively). In this case, 27% of the eGFP-PTS1 diffused freely, while 73%co-diffused with PEX5L. Each dot in the graphs represents an individual FCS measurement. The bars represent the mean and standard deviation. The mean values of the diffusion coefficients of freely diffusing eGFP (dashed red line) and 3xeGFP (fusion protein composed of three eGFP molecules, solid red line) expressed in the same cell line are displayed for comparison with eGFP-PTS1 and PEX5L, respectively.

Specifically, we fluorescently labeled PEX5 with a SNAP-tag (denoted PEX5-SNAP) and the SNAP-tag binding dye Silicon Rhodamine (SiRo, altogether denoted PEX5-SiRo). An artificial cargo protein was created by the conjugation of the tripeptide SKL, a peroxisomal targeting signal type 1, to the C-terminus of the enhanced green fluorescent protein (denoted eGFP-PTS1). We employed a dual expression plasmid to ensure co-expression of both in the same cell (Figure 2b). We started with the longer variant of PEX5, PEX5L-SNAP, and in addition used the point mutant PEX5L S600W-SNAP (or PEX5L S600W-SiRo) as a control, which is not able to interact efficiently with PTS1 cargo proteins such as eGFP-PTS1 (Shimozawa, Zhang et al. 1999)(Stanley et al, Mol.Cell 2006). Using confocal microscopy imaging of eGFP-PTS1, we compared the import efficiency after expressing PEX5L and PEX5L S600W in human fibroblasts, respectively (Figure 2b). Clearly, the wild type (WT) protein showed strong import of eGFP-PTS1 into peroxisomes, as indicated by the higher abundance of fluorescence in the peroxisomes and reduced cytosolic fluorescence background, while cells transfected with PEX5L S600W were characterized by mainly cytosolic fluorescence background. Thus, in the latter case the import was less efficient, which was to be expected since only the pool of endogenous, unlabeled PEX5 could transport and thus import eGFP-PTS1. Figure 2b also highlights representative FCS (or ACF) and FCCS (or CCF) curves of PEX5L-SiRo, PEX5L S600W-SiRo and the co-expressed eGFP-PTS1. While the presence of non-zero and decaying ACFs of all proteins highlights their mobility, the CCFs differ between PEX5L-SiRo and PEX5L S600W-SiRo: The non-zero and decaying CCF of PEX5L-SiRo and eGFP-PTS1 disclose their co-diffusion and thus efficient binding, and the non-existing CCF in the case of PEX5L S600W-SiRo and eGFP-PTS1 reveals their expected missing interaction.

Our FCS study is the first characterization, to our knowledge, of PEX5L and PTS1 mobility in the cytosol of living cells. It is important to mention that we are not able to characterize mobility of PEX5L at the peroxisomal membrane, as the diffusion of PEX5L at peroxisomal membranes is so slow that photobleaching becomes a limiting factor (see Methods section).

### Detailed FCS analysis: Bound and unbound pools of cargo protein PTS1 and slow diffusion of PEX5 independent of PTS1-binding

We next analyzed the FCS data with more detail by fitting equation (1) to the ACF curves and extracting values of the diffusion coefficient D. For the eGFP-PTS1 cargo, we extracted two populations with different mobility, a faster one with D = 48.6 ± 18.6 μm^2^/s that we assigned to unbound eGFP-PTS1, and a slower one which we set to the same value D = 11 μm^2^/s as for PEX5L-SiRo (D = 11.0 ± 3.7 μm^2^/s) to account for eGFP-PTS1 bound to PEX5L-SiRo (see Materials and Methods for details) (Figure 2c). On average, 73% of the eGFP-PTS1 was diffusing bound to the receptor (Figure 2c). However, the fractions of unbound and bound cargo protein varied between independent experiments, probably due to differences in the expression levels. The diffusion coefficient of unbound eGFP-PTS1 (D ≈ 48.6 μm^2^/s) was very close to that of cytosolic eGFP (D = 41.0 ± 11.4 μm^2^/s), which is of similar molecular weight (MW) (26.95 kDa (eGFP) against 27.4 kDa (eGFP-PTS1), Supplementary Table 1) and non-interacting, highlighting free diffusion of unbound eGFP-PTS1 (Figure 2c). Still, these results prove that cargo proteins can be found both bound and unbound to its receptor. As a control, we also studied cells co-expressing the PEX5L S600W mutant and eGFP-PTS1. For PEX5L S600W-SiRo, we found D = 11.9 ± 3.6 μm^2^/s, which is in the same range as PEX5L-SiRo (D = 11.0 ± 3.7 μm^2^/s) and indicates that the mobility of PEX5L was independent of whether PTS1 is bound (PEX5L-SiRo) or not (PEX5L S600W-SiRo). Further, we again could fit the FCS data of eGFP-PTS1 with two components, the unbound free form (D = 48.6 ± 18.6 μm^2^/s, as before), and a bound form with D = 11.0 μm^2^/s set to the mobility of PEX5L. However, the latter fraction was much lower (on average 12%) than for the previous wild-type PEX5L-SiRo expressing cells (on average 73 %), since now only a small number of endogenous fully functional PEX5 was available to bind eGFP-PTS1 (Figure 2c).

In both wild-type PEX5L-SiRo and PEX5L S600W-SiRo expressing cells, the respective FCS data were both well described by a single diffusing population (D ≈ 11 μm^2^/s). This finding was somehow surprising, as two populations might have been expected, one free and one carrying PTS1 cargo proteins. However, as highlighted already in the previous paragraph, the mobility of PEX5L seemed to be independent of whether PTS1 was bound or not. To further detail this, we compared the mobility of PEX5L-SiRo and PEX5L S600W-SiRo against the similar weighed aggregate of three eGFP molecules (80.85 kDa for 3xeGFP compared to 91.39 kDa for PEX5L-SNAP, Supplementary Table 1). 3xeGFP showed a diffusion coefficient of about twice as high (21.9 ± 4.9 μm^2^/s) as PEX5L, i.e. double the mobility (Figure 2c). Also, we confirmed the general slow and PTS1-independent diffusion of PEX5L for other cargos such as eGFP-catalase and eGFP-SCP2. In all cases, diffusion of PEX5L-SiRo was equally slow (Figure 3a). Consequently, the slow-down in diffusion of PEX5L is not due to its molecular weight and independent of PTS1 cargo binding.

**Figure 3:**
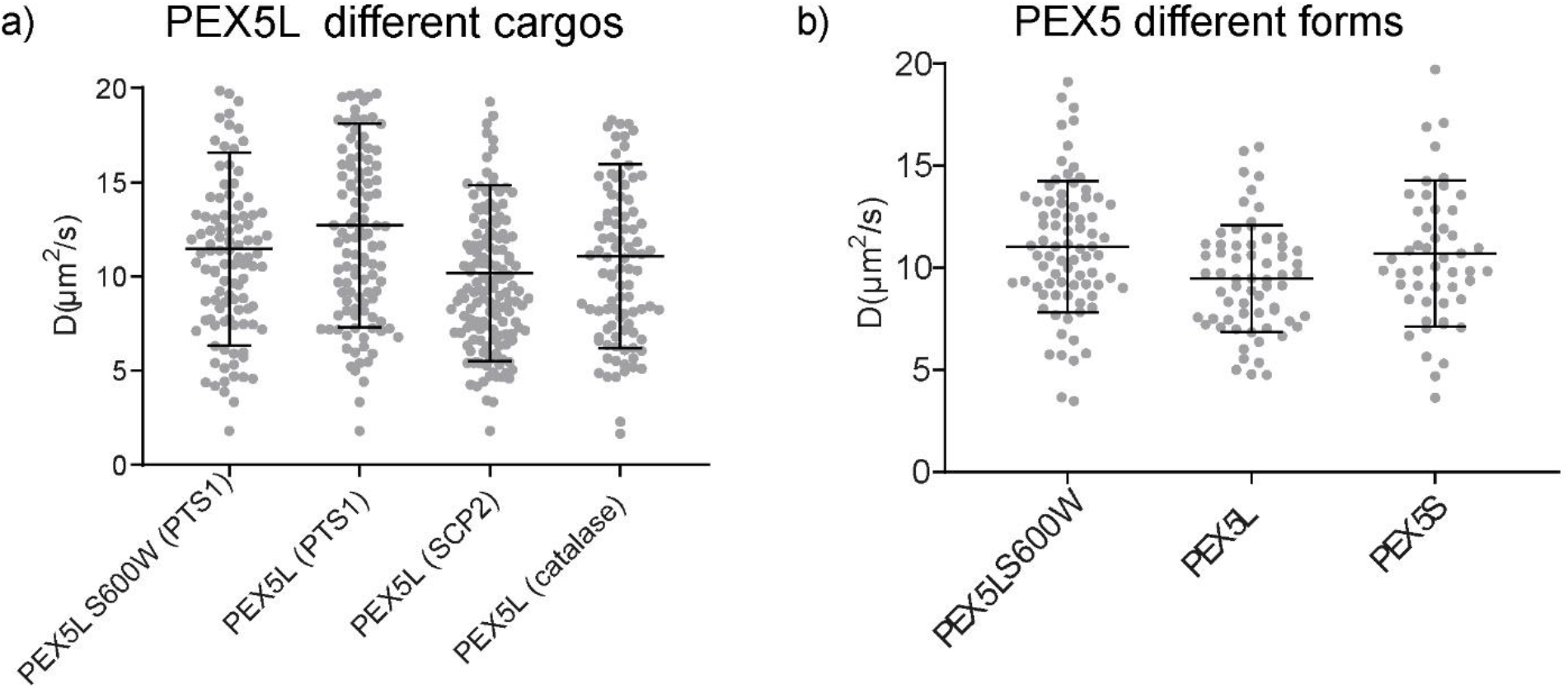
The diffusion of PEX5 is not influenced by binding cargo proteins or the PTS2 pathway. a): The diffusion coefficient of PEX5L is independent of its cargo protein. Human fibroblasts were transfected with the dual plasmids PEX5L S600W/eGFP-PTS1, PEX5L/eGFP-PTS1 or with two plasmids one encoding PEX5L-SNAP and the other eGFP-SCP2 or eGFP-Catalase. The diffusion coefficient of PEX5L did not change upon co-expression of different cargo proteins. b): Diffusion coefficients of PEX5L and PEX5S. Human fibroblasts were transfected with PEX5L S600W, PEX5 long isoform (PEX5L) and PEX5 short isoform (PEX5S) respectively, all on dual expression plasmids with eGFP-PTS1. PEX5L has an additional binding site for the PTS2 receptor protein PEX7, which is lacking in PEX5S. As the diffusion coefficients of the different forms of PEX5 did not significantly differ, an influence of the PTS2 pathway on the diffusion coefficient of PEX5 was excluded. Each dot in the graph represents an individual FCS measurement, performed on three independent biological replicates. The bars represent the mean and standard deviation.

### The PTS2 pathway has no influence on PEX5 diffusion

As highlighted before, the PEX5 protein exists in two different isoforms, a long PEX5L and a short PEX5S form. While both isoforms interact with PTS1 cargo proteins, the long isoform PEX5L also contains a binding site for the PTS2-receptor PEX7. Thus, cargo-loaded PEX7 binds to PEX5L and bridges the binding of PTS2 cargo proteins to the PTS1 import receptor PEX5. To investigate any influence of the binding of PEX7/PTS2 complexes on the mobility of PEX5, we compared the diffusion of the PEX5L and PEX5S isoforms. Both variants were again jointly expressed in human fibroblasts (GM5756-T) from dual PEX5L-SNAP/eGFP-PTS1 and PEX5S-Halo/eGFP-PTS1 plasmids, respectively, and fluorescent labeling realized with SiRo-SNAP and SiRo-Halo, respectively. Again, both variants showed undistinguishable mobility (Figure 3b), and we concluded that binding of PEX7 and its cargo was not rate-limiting for the cytosolic mobility of PEX5.

### Super-resolution study: PEX5L diffuses freely

As PEX5L showed an unexpectedly slow diffusion in the cytosol, we wanted to explore any heterogeneity in mobility (e.g. due to transient interactions) by comparing its diffusion mode with a free diffusing molecule (GFP-SNAP). For this, we applied FCS on a super-resolution STED microscope, optimized with adaptive optics (AO-z-STED-FCS) (Barbotin et al., 2019b). STED-FCS allows to measure the mobility of a fluorescent molecule while changing the size of the observation volume from around 200-250 nm and 700 nm in lateral and axial diameter to below 50-80 nm and 300 nm, respectively, providing otherwise inaccessible information on molecular diffusion dynamics. A prominent example is the distinguishing of free and hindered diffusion modes such as due to transient, interaction-evoked slow-downs (Eggeling et al., 2009a). While STED-FCS is an established technique to study diffusion dynamics in two dimensions such as on membranes (Sezgin et al., 2019), our application at a three dimensional (3D) cytosolic level required refined technical implementation (Gao et al., 2017; Ringemann et al., 2009). In order to tune the size of the effective fluorescence observation spot along the axial z-direction, a top-hat intensity-shaped STED laser beam was overlapped with the standard excitation beam, whereby the performance was optimized by reducing possible optical aberrations using adaptive optics (AO) (Barbotin et al., 2019a). In the final AO-z-STED-FCS measurements, the STED laser power was stepwise increased to record FCS data and thus determine cytosolic PEX5L mobility at varying observation volumes (Figure 4a). We first characterized the axial confinement of our observation volume from a standard confocal volume to the maximum compressed volume by acquiring a series of images of 40-nm sized fluorescent beads at different STED laser powers, highlighting a reduction of the axial diameter from 671 nm down to 256 nm (Figure 4b). We then used AO z-STED-FCS to measure the diffusion of eGFP-SNAP (labeled with SiRo dye) in the cytosol of living cells. eGFP-SNAP is an artificial protein that is (to our knowledge) not interacting with any cellular component and is therefore a suitable control to represent free diffusing modality. Its transit times through the observation volume decreased in coincide with increasing STED power and thus confinement (Figure 4c), indicating the absence of hindered diffusion (Ringemann et al., 2009). Using AO z-STED-FCS, we also highlighted free diffusion for cytosolic PEX5L and PEX5L S600W (labelled with SiRo). For both PEX5L and PEX5L S600W the decrease in transit time with confinement of observation volume followed the same pattern as for freely diffusing eGFP-SNAP fusion protein (Figure 4c). Consequently, the slowed down diffusion of PEX5L in the cytosol is not due to transient interactions with other more immobilized binding partners, but rather a stable interaction with a permanent partner. Notably, this interaction is independent of the ability of PEX5 to bind to cargo proteins (as highlighted for non PTS1-binding PEX5L S600W).

**Figure 4:**
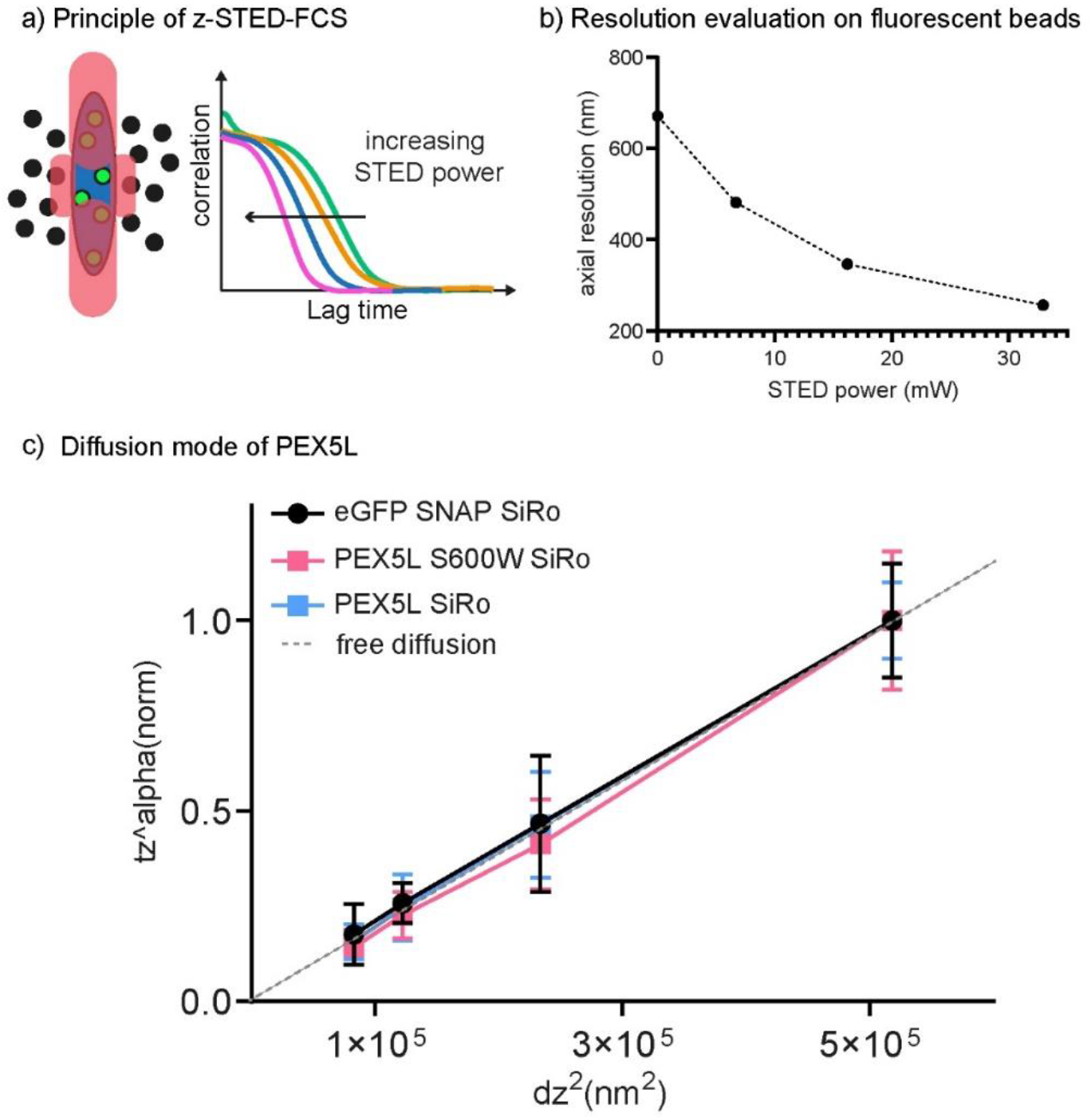
PEX5L diffuses freely, as shown by AO-z-STED-FCS. a): Principle of z-STED-FCS. STED-FCS experiments determine the apparent diffusion coefficient from FCS measurements at different observation spots sizes. The effective size of the observation spot (blue) decreases by increasing the power of the z depletion STED beam (red, left), which shortens the decay time of the ACFs (right). By analysing the transit times in dependence of the observation area, mobility information at the sub-diffraction level can be extracted. b): Resolution calibration for z-STED on fluorescent beads. To calibrate the resolution for the z-STED depletion beam 40 nm crimson beads were imaged at different STED powers (0 mW, 7 mW, 16 mW, 33 mW measured at the back aperture of the objective lens) and the resolution was evaluated. As the STED power increases the resolution along the optical axis increases accordingly (671 nm, 481 nm, 346 nm, 256 nm). c): AO zSTED-FCS measurements of PEX5L diffusion. PEX5L, PEX5L S600W and the eGFP-SNAP fusion protein used as control for free diffusion (all labelled with SiRo) were expressed in human fibroblasts. The diffusion of these proteins was measured with z-STED FCS combined with AO, to correct for aberration, at different STED laser powers and therefore different sizes of the observation spot. The relative decrease in axial transit times^alpha (fixed at t_z_^0.75), normalised to confocal, was plotted against the square of the observation focus axial size (dz^2^). Both quantities are proportional in presence of free diffusion (equation 1). PEX5L and PEX5L S600W showed the same diffusion behaviour as the eGFP-SNAP fusion protein as well as the free diffusing trend line represented on the graph (grey dotted line), disclosing a free diffusion of PEX5. STED powers were tuned at 0 mW, 7 mW, 16 mW, 33 mW. Each dot in the graph represents the mean value of at least 30 measurements at each STED power, performed on three independent biological replicates. Bars represent the mean and standard deviation.

### PEX5L does not form homo-oligomers within the cytosol

As highlighted, PEX5L integrates into the peroxisomal membrane to guide cargo proteins into the peroxisomal matrix. Due to its accumulation at the membrane, it was previously hypothesized that PEX5L might oligomerize at the membrane or even already in the cytosol(Gould and Collins, 2002). To investigate if the observed slow cytosolic diffusion of PEX5L is linked to a possible homo-oligomerization, we conducted a FCCS study between differently labeled PEX5L to highlight their possible co-diffusion and thus potential homo-oligomerization. Here, cells expressing PEX5L-SNAP were incubated with a solution containing a 1:1 (mol:mol) mixture of the red-fluorescing SNAP-SiRo and green-fluorescing SNAP-Cell505 dyes. While the autocorrelation curves for each signal (SiRo and Cell505) were quite similar (Figure 5b) and confirmed the slow diffusion (10.4 ± 4.4 μm^2^/s and 12.8 ± 4.2 μm^2^/s for SiRo and Cell505 labelled PEX5L, respectively), as expected for the similarly tagged PEX5L proteins, there was no evidence of any cross-correlation signal. While we cannot exclude a weak or very transient interaction, we did not find any evidence of a strong PEX5L homo-oligomerization. In addition, the cytosolic mobility of PEX5L did also not change in a CRISPR/Cas9 derived PEX5 knock out (KO) cell line (Supplementary Figure 2), precluding oligomerization of PEX5L with endogenous unlabelled PEX5.

**Figure 5:**
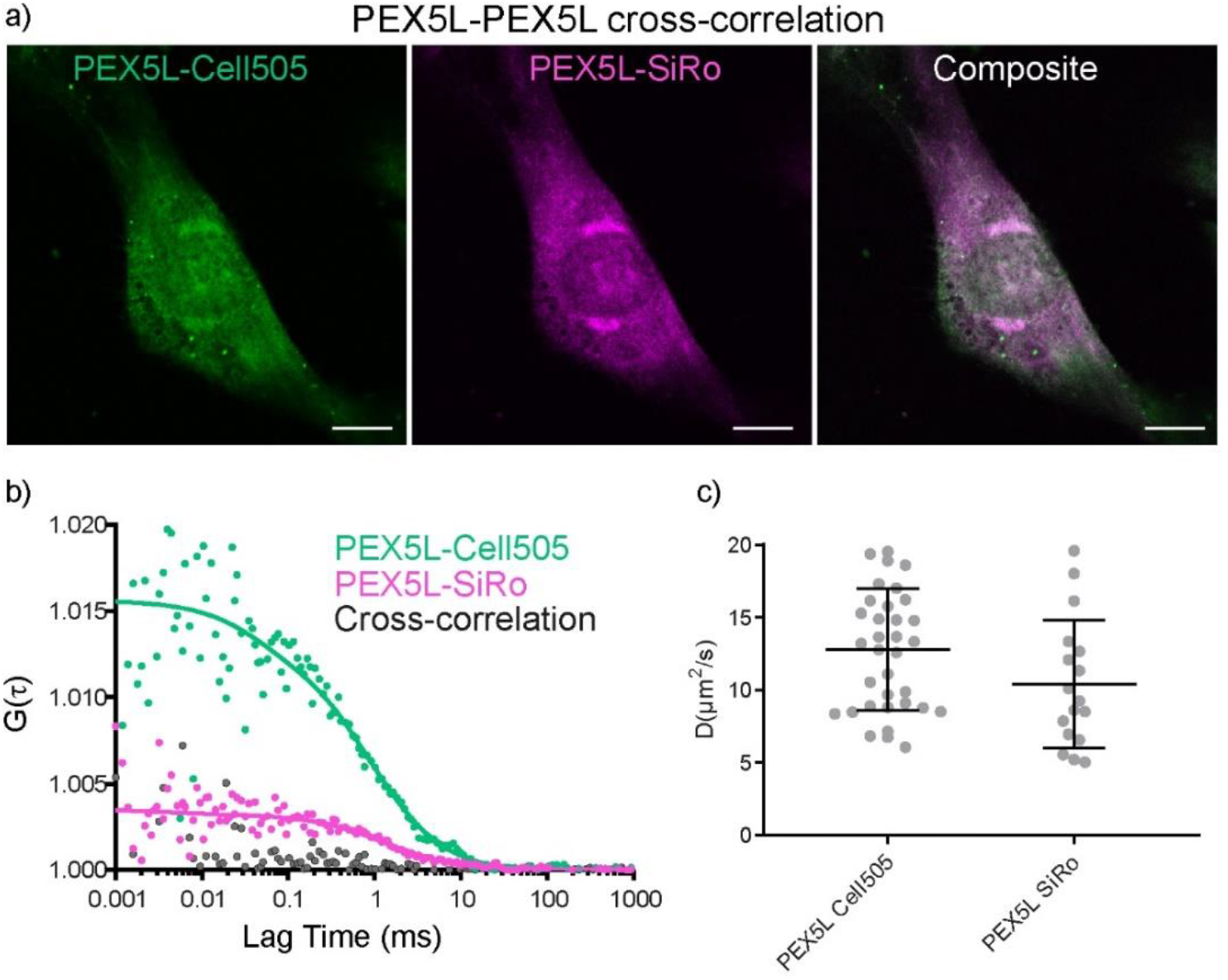
PEX5L does not form homo-oligomers in the cytosol. a): Human fibroblasts expressing PEX5L-SNAP were co-incubated with SNAP-Cell505 and SNAP-SiRo dyes in a 1:1 (mol:mol) ratio. The protein was mainly localized in the cytosol. Scale bar 10 μm. b): Representative autocorrelation curves of PEX5L-SNAP labelled with SNAP-Cell505 (green) and SNAP-SiRo (magenta) are displayed. No cross-correlation signal between the differently labelled PEX5L proteins could be measured, indicating no abundant interaction between PEX5L molecules in the cytosol. c): The diffusion coefficient of PEX5L-SNAP-Cell505 and PEX5L-SNAP SiRo did not differ. Each dot in the graph represents an individual FCS measurement. The bars represent the mean and standard deviation.

### Peroxisomal membranes and the cytoskeleton have no influence on the diffusion of PEX5L

PEX5L interacts with peroxisomal membrane proteins and integrates into the membrane during cargo-translocation. Such interaction with the peroxisomal membrane or membrane proteins could be a cause of the observed slow-down. Also, there are indications that PEX5L interacts with other organelles like liposomes (Kong et al., 2020). Here, two different approaches were taken to test if an interaction with peroxisomal membranes or other organelles could explain the slow PEX5L diffusion.

First, a cell-free model system in form of giant plasma membrane vesicles (GPMVs) was used. Here, upon treatment of the cells with NEM (N-Ethylmaleimide), the plasma membrane gets detached from the cytoskeleton and forms free-standing vesicles that contain cytosolic proteins and are devoid of organelles and cytoskeleton (including microtubules and actin filaments) (schematic Figure 6a) (Schneider et al., 2017; Sezgin et al., 2012). Therefore, proteins in these vesicles cannot interact with the cytoskeleton or intracellular membranes. To compare the diffusion of PEX5L in cells and in GPMVs, we expressed PEX5L-eGFP (MW 99.3 kDa) and 3xeGFP (MW 80.85 kDa) in cells, generated GMPVs by treatment with NEM, and determined the diffusion coefficient of PEX5L-eGFP and 3xeGFP in both systems using FCS as before: D = 10.3 ± 2.6 μm^2^/s (cells) and 19.8 ± 10.8 μm^2^/s (GPMVs) for PEX5L-eGFP, and D = 25.0 ± 4.2 μm^2^/s (cells) and 38.9 ± 13.5 μm^2^/s (GPMVs) for 3xeGFP (Figure 6b). For both PEX5L-eGFP and 3xeGFP there was a general increase in mobility in GPMVs compared to cells (factor 1.9 for PEX5L and 1.6 for 3xeGFP), owing to the decreased cytosolic crowding in GPMVs (Schneider et al., 2017; Sezgin et al., 2012). Most importantly, the difference in mobility, or ratio between diffusion coefficients D, was similar in cells and GPMVs (D(3xeGFP)/D(PEX5L-eGFP) = 2.4 in cells compared to 2.0 in GPMVs), highlighting a slow-down of cytosolic PEX5L independent of potential interactions with organellar membranes or the cytoskeleton.

**Figure 6:**
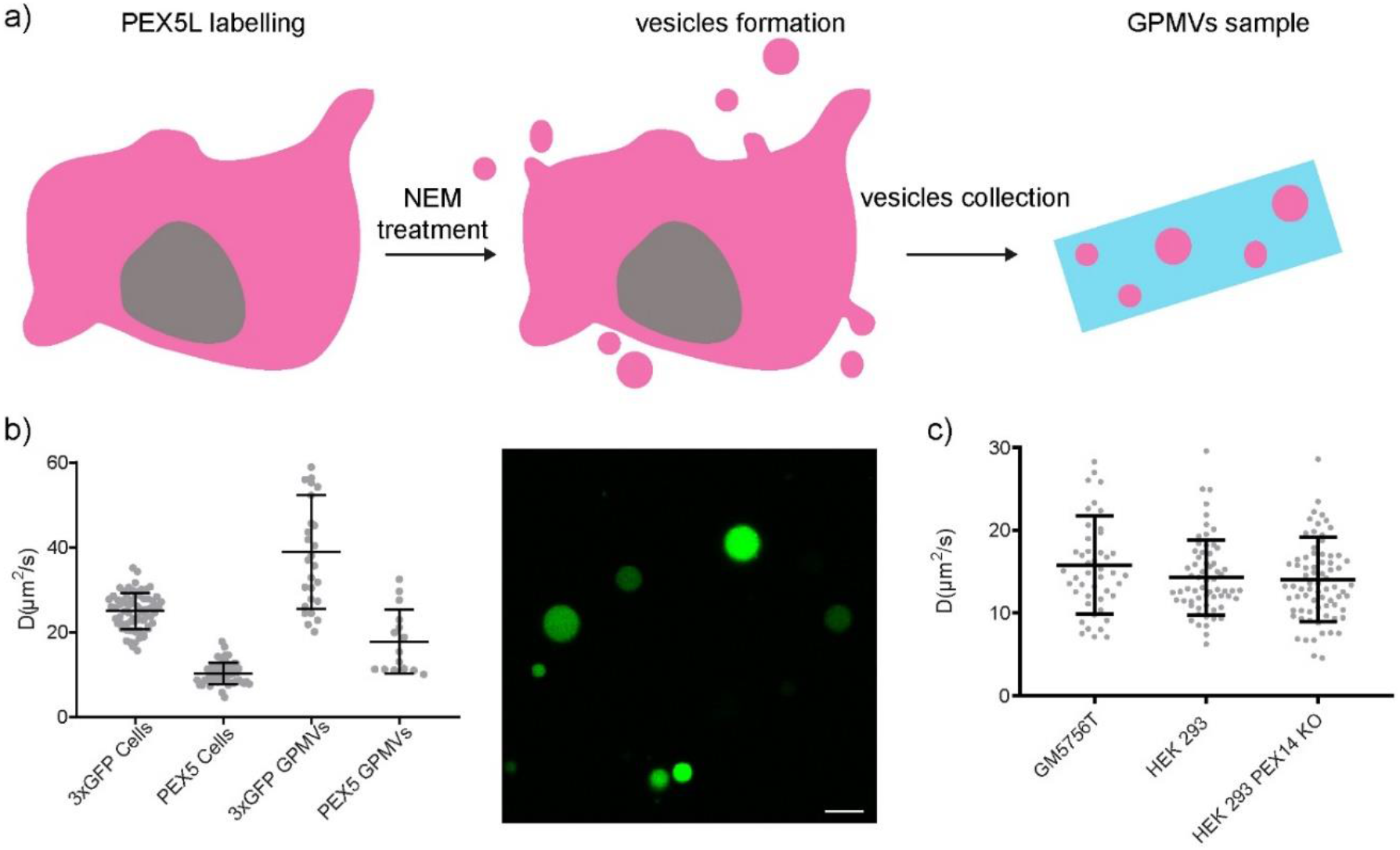
PEX5L is not slowed down by peroxisomal membranes or cytoskeleton. a): Principle of GPMV formation. Cells expressing the labelled proteins of interest were incubated with NEM, which disrupts the interaction between the cortical actin and the plasma membrane, inducing the formation of blebs of the plasma membrane. These blebs contain cytosolic proteins but no organelles or a filamentous cytoskeleton. After blebbing of from the cells the GPMVs can be collected and imaged. b): Diffusion coefficient measured in intact cells and GPMVs. Human fibroblasts were transfected either with 3xeGFP, which has a molecular weight of 81 kDa, close to that of PEX5L-eGFP (99.3 kDa), or PEX5L-eGFP itself. The diffusion coefficients were measured directly in these cells, or in GPMVs derived from these cells, containing cytosolic proteins only. Although the diffusion of the 3xeGFP and PEX5-eGFP became faster in the GPMVs as expected (Schneider et al., 2017), the ratio of the two different proteins diffusion remained comparable (2.42 in cells and 1.96 in GPMVs). Each dot in the graph represents an individual FCS measurement, performed on three independent biological replicates. On the right, a microscopic image of the measured GPMVs containing eGFP labelled proteins is shown. Scale bar 10 μm c): The cytosolic diffusion of PEX5L is independent of PEX14. PEX5L was expressed together with eGFP-PTS1 in human fibroblasts (GM 5756T), HEK 293 cells and the HEK KOPEX14 cell line. The diffusion speed of PEX5L was measured in WT fibroblasts (15.8 ± 5.9 μm^2^/s), WT HEK cells (14.3 ± 4.5 μm^2^/s) and WT HEK KO PEX14 cells (14.0 ± 5.0 μm^2^/s). As the diffusion coefficients are comparable in all three tested cell lines an influence of PEX14 on the diffusion of PEX5L can be excluded. Each dot in the graph represents an individual FCS measurement, performed on three independent biological replicates. The bars represent the mean and standard deviation.

In a second approach, we investigated a potential interaction of PEX5L with peroxisomal membranes by determination of PEX5L mobility in a PEX14-deficient knockout (KO) cell line. As PEX5L binds PEX14 at the peroxisomal membrane (Will et al., 1999), the interaction of PEX5L with peroxisomal membranes should be inhibited in the absence of PEX14 or at least significantly decreased. Here, the dual plasmid expressing PEX5L and eGFP-PTS1 was expressed in a HEK 293 PEX14 KO cell line created by CRISPR/Cas9. In these cells, the diffusion coefficient of PEX5L (D = 14.0 ± 5.0 μm^2^/s) was however comparable to that in WT fibroblasts (15.8 ± 5.9 μm^2^/s) or WT HEK cells (14.3 ± 4.5 μm^2^/s), indicating that the interaction of PEX5L with PEX14 does not cause its cytosolic slow down (Figure 6c).

### Diffusion of PEX5L N- and C-terminal halves

PEX5L can be divided into two different functional parts, the structurally disordered N-terminal half (amino acids 1-335) (Costa-Rodrigues et al., 2005; Shiozawa et al., 2009), containing several WxxxF/Y motives that play a role in docking to the peroxisomal membrane (Freitas et al., 2011; Neuhaus et al., 2014), and the globular C-terminal half (amino acids 314-639) containing tetratricopeptide repeat (TPR) motifs, which interact with the PTS1 signal sequence (Saidowsky et al., 2001) (schematic in Figure 1, Figure 7a top)

**Figure 7:**
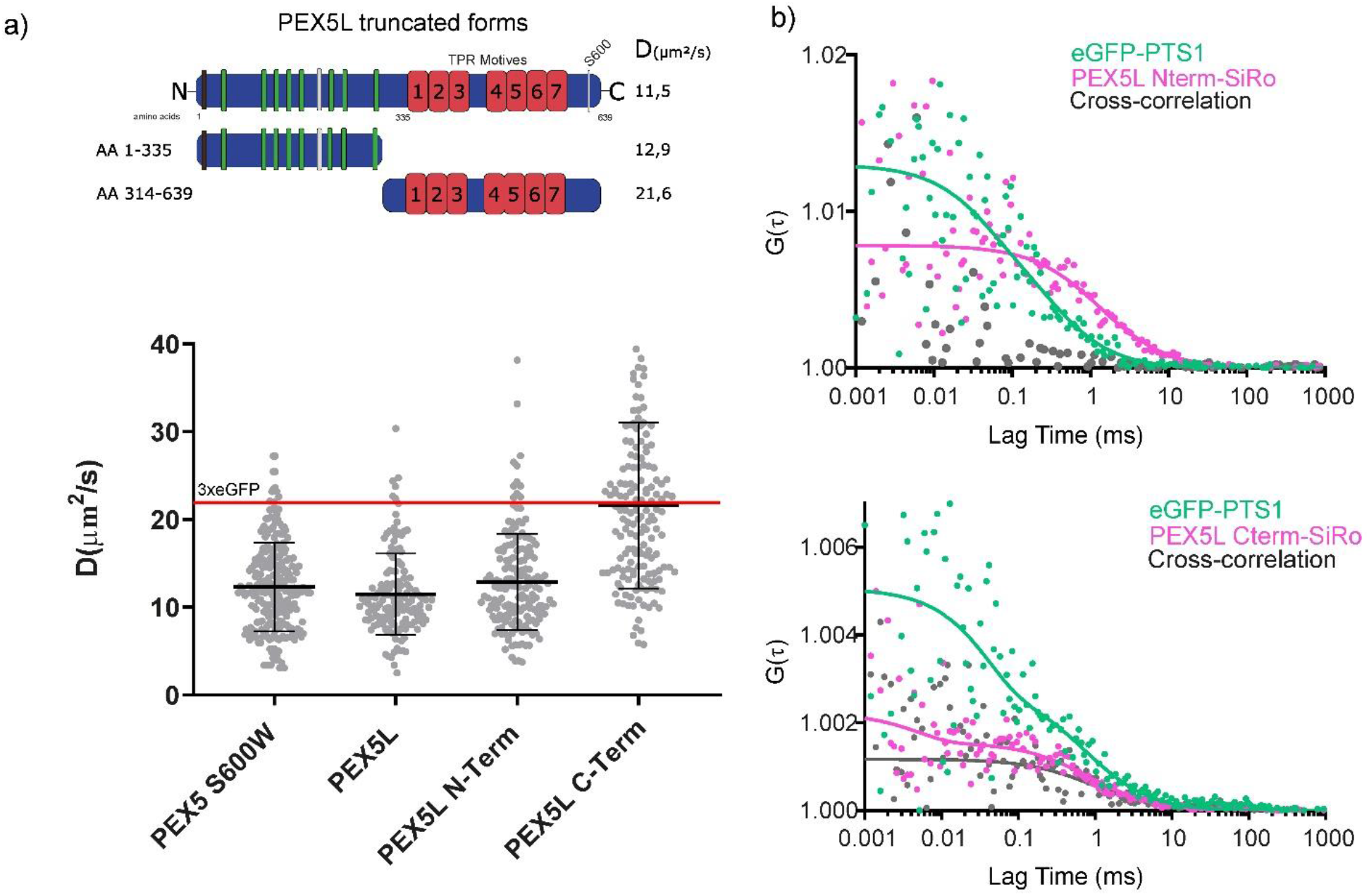
The N-terminal part of PEX5 slows down its diffusion. a): Schematic representation of PEX5L. PEX5L can be divided into two different functional parts, the unstructured N-Terminus (AA 1-335) with the WxxxxF/Y (green) motives and the globular C-Terminus (AA 314-639) with 7 TPR motives (red), which are interacting with the PTS1 proteins. The PEX7 binding site is shown in grey and the ubiquitin binding site in black at the very C-terminus. Truncated versions of the full length PEX5L were created to analyse the diffusion of the two functional parts of PEX5L separately. PEX5L S600W (which cannot interact with PTS1 cargo proteins), PEX5L, PEX5L (1-335) (N-Term) and PEX5 (314-639) (C-Term) were expressed separately in human fibroblasts and their diffusion coefficients were measured using FCS. The C-terminus diffused at 21.6 ± 9.4 μm^2^/s, while the N-terminus at 12.9 ± 5.5 μm^2^/s, similarly as the full-length PEX5L (11.5 ± 4.6 μm^2^/s), indicating that this part defines the diffusion speed of the full-length protein. Each dot in the graph represents an individual FCS measurement, performed on three independent biological replicates. The bars represent the mean and standard deviation. b): The autocorrelation functions of eGFP-PTS1 (green) and PEX5L N-terminus (upper panel) or PEX5L-C-terminus (lower panel) (magenta) are displayed. The C-terminus of PEX5L diffused together with the eGFP-PTS1 cargo protein, as a cross-correlation could be measured, while the N-terminus of PEX5L did not diffuse together with the cargo protein. Here, no cross-correlation could be detected.

We created truncations of PEX5 that comprise either the N-terminal or the C-terminal half of PEX5L (referred to as PEX5L N-Term and PEX5L C-Term), expressed them together with eGFP-PTS1 (in dual expression plasmids) in human fibroblasts, and determined their diffusion coefficients as well as co-diffusion with eGFP-PTS1 using FCS and FCCS as described before. As expected, FCCS highlighted co-diffusion and thus binding between PEX5L C-Term and eGFP-PTS1, while PEX5L N-Term did not (Figure 7b). On the other hand, mobility of PEX5L N-Term was similarly slow (D = 12.9 ± 5.5 μm^2^/s) as full length PEX5L (11.5 ± 4.6 μm^2^/s) and PEX5L S600W (12.32 ± 5.1 μm^2^/s), while diffusion of PEX5L C-Term was two-fold faster (D = 21.6 ± 9.4 μm^2^/s) and about the same as 3xeGFP (D = 21.9 ± 4.9 μm^2^/s) (Figure 7a). Interestingly, PEX5L C-Term and PEX5L N-Term have similar molecular weight (58 kDa and 56.5 kDa, respectively). Therefore, we concluded that the slowing-down element of the cargo receptor PEX5 is located in its N-terminal half.

### The slow diffusion of PEX5L-N-terminal half is caused by a cytosolic factor

The finding that the N-terminal half of PEX5L is responsible for the slow diffusion of the protein raised the question whether this slow diffusion was caused by a specific cytosolic N-terminal binding partner or by the structurally disordered nature of the N-terminal half compared to the more ordered C-terminal PTS1-binding domain. Therefore, we compared the mobility of PEX5L N-Term with that of a similarly unstructured protein, the N-terminal half of PEX5 from *Trypanosoma brucei*. While the structural architecture of PEX5L (and therefore also of its N-terminus) is similar between trypanosomes and the human PEX5L protein (Figure 8a), we expected cytosolic interactions to differ between both variants due to their descent. Therefore, we expressed the N-terminal half of human PEX5L (*Hs*PEX5L N-Term, AA 1-335) and the N-terminal half of PEX5 from trypanosoma (*Tb*PEX5 N-Term, AA 1-340) in human fibroblast cells (labelled via SiRo-SNAP as before). Here, *Tb*PEX5 N-Term diffused almost twice as fast as its human counterpart (Figure 8c). To test whether this effect was caused by a cytosolic interaction partner of the human PEX5L, we created recombinant versions of the two proteins, both fused to eGFP. This allowed us to measure the diffusion of the proteins in solution, i.e. without the presence of any potential binding partner. Here, both *Hs*PEX5L N-Term and *Tb*PEX5 N-Term diffused similarly fast (Figure 8d). Consequently, the bulky unstructured character of the N-terminal part of PEX5 was not the main reason for the slow-down, but it was rather a human-specific cytosolic binding partner. To test the latter, we isolated cytosolic components from HEK 293 cells and incubated them with the recombinant proteins *Hs*PEX5L N-Term and *Tb*PEX5 N-Term (Figure 8b). Interestingly, the human-based cytosolic components specifically slowed down human PEX5L N-Term but not its trypanosomal counterpart (Figure 8d), indicating the interaction with a cytosolic factor as the main cause of slow-down in diffusion.

**Figure 8:**
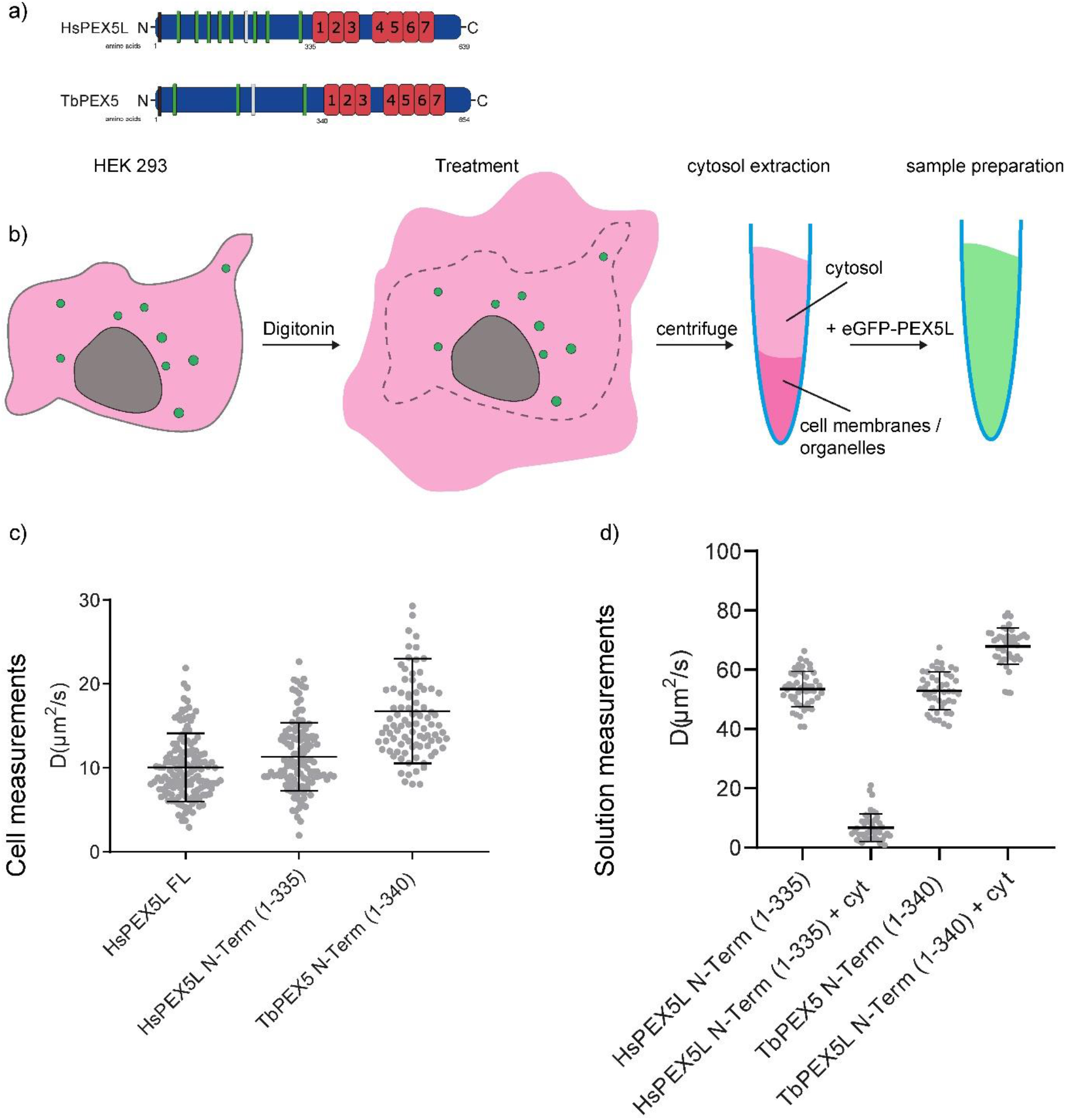
The slow diffusion of human PEX5 is caused by a cytosolic factor. a): Architectural similarity of human and trypanosomal PEX5. The PTS1-receptor PEX5 from human and trypanosomes are similar in size and contain homologous structures as the C-terminal TPR motives (red) which bind PTS1-cargo proteins, and the WxxxxF motives in the N-terminal half (green), which mediate the binding of PEX5 to the peroxisomal membrane proteins PEX13 and PEX14. Also, both proteins contain a PEX7 binding site (grey) and an ubiquitin binding site (black) at the very N-terminus. b): Extraction of the cytosol from human HEK 293 cells. Untreated HEK 293 cells were harvested and treated with a digitonin containing buffer. The digitonin is exclusively permeabilizing the plasma membrane of the cells, so the cytosolic content is set free. After centrifugation the cell membranes and organelles remain in the pellet, while cytosolic proteins can be found in the supernatant. This supernatant gets mixed with the recombinant eGFP labelled proteins and FCS is performed (result in D). c): Diffusion coefficients in cells. Human fibroblasts were transfected with plasmids encoding for PEX5L, the N-terminal half of human PEX5L (*Hs*PEX5L N-Term, AA 1-335) and the N-terminal half of PEX5 from *Trypanosoma brucei* (*Tb*PEX5 N-Term, AA1-340), and labelled with SNAP-SiRo. The diffusion coefficient of the human protein is significantly lower than the diffusion coefficient of the trypanosomal PEX5. Each dot in the graph represents an individual FCS measurement, performed on three independent biological replicates. The bars represent the mean and standard deviation. d): Diffusion coefficients in solution. *Hs*PEX5 N-Term-eGFP and *Tb*PEX5 N-Term-eGFP were recombinantly expressed in *E. coli* and purified. The diffusion coefficients of the human and trypanosomal N-terminal halves of PEX5 did not differ. The recombinant proteins were mixed with the cytosol isolated from human HEK cells. This led to a slowdown of the human PEX5L N-terminal half but not for the trypanosomal *Tb*PEX5 N-Term-eGFP. Each dot in the graph represents an individual FCS measurement, performed on three independent biological replicates. The bars represent the mean and standard deviation.

## Discussion

In this study we applied advanced microscopy and spectroscopy techniques combined with biological methods to investigate the diffusion behavior of the peroxisomal import receptor PEX5 and its peroxisomal cargo proteins in the cellular cytosol. The combinatory approach revealed a broad range of biophysical information on cargo recognition and the migration behavior of the free and cargo-loaded receptor in the cytosol.

Using FCS, we could fully characterize the diffusion properties of PEX5 and we could reveal and characterize features expected from its function in peroxisomal protein import: (1) Co-diffusion in a complex with cargo proteins, proved by the cross-correlation of PEX5 and PTS1 (Figure 2b). (2) Diffusion of the major fraction of PTS1 in a complex with PEX5 (73%), and only a minor part with free diffusion characteristics, i.e. not bound to PEX5 (27%), but this distribution might be influenced by the overexpression of the proteins in this experimental setup (3) By cross-correlation analysis employing C- and N-terminal truncations of PEX5, we could demonstrate in living cells that the C-terminal part but not the N-terminal part of PEX5 binds PTS1-cargo proteins (Figure 7b) which formerly has only be seen in vitro (Harano et al., 2001). (4) Using super-resolution AO z-STED-FCS, we could prove cytosolic free diffusion of PEX5 (Figure 4c).

Strikingly, the analysis of the diffusion characteristics of PEX5 also revealed a very slow cytosolic diffusion of PEX5, which was much slower than expected from its molecular weight. Further, this slow diffusion was independent of (1) binding of PTS1-cargo (Figure 2) and cargo type (Figure 3a), (2) interaction with PTS2-cargo and its cargo receptor PEX7 (Figure 3b), (3) possible (transient) interactions with peroxisomal membranes and other organells (using PEX14 knock-out cells and GPMVs) (Figure 6c), (4) cytoskeleton meshwork (Figure 6b), (5) the structurally disordered N-terminal half of PEX5 (Figure 7 and 8c), and (6) possible and non-confirmed cytosolic oligomerization (Figure 5). Related to the latter issue, PEX5 binding of oligomerized cargo proteins has been reported before as it is a precondition of piggy-back transport of proteins into peroxisomes (Glover et al, 1994; MyNew and Goodman, 1994). Along this line, a dimeric alanine-glyoxylate amino-transferase (AGT) could also bind two PEX5 receptors (Fodor et al., 2015b). However, it is still a matter of debate whether PEX5 binds oligomerized cargos, which then can form large complexes with several receptor proteins involved. This “preimplex” hypothesis was originally described by Gould and Collins (Gould and Collins, 2002) and supported by findings that peroxisomal enzymes enter large protein complexes before they get translocated into the peroxisomal matrix (Bellion and Goodman, 1987). Although we cannot exclude possible very short and transient interactions, our data indicated no presence of PEX5 oligomerization in the cytosol, which is in line with other previous studies (Costa-Rodrigues et al., 2005).

Finally, we compared cytosolic diffusion of human PEX5L with that of PEX5 from *Trypanosoma brucei* (Sampathkumar et al., 2008), disclosing distinct mobility differences. From the fact that *Tb*PEX5 diffused faster than its human counterpart in the cytosol of mammalian cells, we discovered (1) that the unstructured N-terminal of PEX5 is not the cause for the non-typical slow diffusion of the receptor but that it only has a minimal influence on its mobility (as both variants are characterized by such an unstructured part), and (2) that the slow diffusion of PEX5 is not caused by the structure of the protein but by binding to another cytosolic protein. This was confirmed by *in vitro* measurements on recombinant human PEX5 and *Tb*PEX5. While both recombinant variants of PEX5 showed no difference in their diffusion behaviour in solution, only the human PEX5 was slowed down in the presence of cytosolic proteins. In addition, diffusion of PEX5L was very distinct slow rather than heterogeneous over a larger range of mobilities, i.e. the interaction rather stable and non-transient. These findings point to a so-far unknown cytosolic interaction partner that binds to the N-terminal part of human PEX5 and determines its peculiar diffusion behavior. The identity of this interaction partner still remains to be shown. Possible are interactions with a larger chaperon assembly, which would accompany PEX5 in its recognition of cargo proteins or protect the intrinsically disorder N-terminal region of PEX5 from aggregation. Also, an interaction of PEX5 with ribosomes could be envisioned, which would be in line with the observation that the mRNAs for the synthesis of peroxisomal proteins are found in close proximity to peroxisomes (Zipor et al, 2009).

Besides novel insights into diffusion and interaction dynamics of peroxisomal proteins and especially the essential cargo-carrier and -import protein PEX5 in the cytosol of living cells, our study highlights the potential of using complementary experimental tools from advanced fluorescence microscopy and spectroscopy over model systems to biochemical and molecular biology approaches to decipher molecular interactions in the cytosol via studying their diffusion dynamics. Here, the combinatory approach revealed characteristics of the cytosolic migration behavior of peroxisomal proteins, their receptor interaction prior to peroxixomal targeting and import and it disclosed the cytosolic interaction of the peroxisomal import receptor PEX5 with a novel high-molecular weight binding partner.

## Materials and Methods

### Plasmids

Sequences of the primers used are shown in Supplementary Table 2.

For the simultaneous expression of different PEX5 variants together with eGFP-SKL (eGFP-PTS1), we used the dual expression plasmid pIRES2/eGFP-SKL as described before (Neuhaus et al., 2014). First, a SNAP-tag was integrated by amplifying the SNAP sequence with plasmids RE4692/RE4693 and restriction sites SalI/BamHI. Afterwards the full-length PEX5L was amplified from pIRES2 PEX5L/eGFP-SKL (Fodor et al., 2015a) with primers RE4640/RE4641 and cloned into the pIRES2 SNAP/eGFP-SKL using BglII/SalI restriction sites. This pIRES2 PEX5L-SNAP/eGFP-SKL plasmid was used to construct all variations of PEX5L for the FCS measurements. Most of these variations were created using FastCloning (Li et al., 2011). In brief, the vector backbone and the insert were amplified by PCR with overlapping ends.

Afterwards, PCR products were mixed, the template DNA was digested with DpnI and overlapping sticky ends were annealed. Here, the vector backbone was amplified from the pIRES2 PEX5L-SNAP/eGFP-SKL and the insert from different sources: PEX5-C-Term (pIRES2 PEX5L 1-335-SNAP/eGFP-SKL): Vector amplification with primers RE4816/ RE4817; and insert amplification from pIRES2 PEX5L/eGFP-SKL with primers RE6194/RE6195.

For PEX5-N-Term (pIRES2 PEX5L 314-639-SNAP/eGFP-SKL) the PEX5L fragment was amplified with the primers RE6196/RE6197 and subcloned into BglII/SalI digested pIRES2 PES5L-SNAP/eGFP-SKL.

For PEX5 S600W-SNAPeGFP-SKL the PEX5 S600W sequence was amplified with the primers KR001/KR002 from PEX5 S600W (Shimozawa et al., 1999) in pcDNA3.1 and ligated into the BgIII/SalI digested pIRES2 PEX5-SNAP/eGFP-SKL.

The PEX5S-HALO/eGFP-SKL plasmid was created from the pIRES2/eGFP-SKL as well. First the HaloTag was integrated by amplification with primers RE4694/RE4695 and subcloning of the PCR-product into the pIRES2/eGFP-SKL using restriction sites SalI/BamHI. Afterwards, the full-length PEX5S was amplified with primers RE4640/RE4641 and cloned into the pIRES2 HALO/eGFP-SKL using BglII/SalI restriction sites.

pIRES2 *Tb*Pex5 1-340 SNAP eGFP-SKL: Vector amplification with primers RE6496/RE6486;Insert amplification from *Tb*PEX5 (De Walque et al., 1999) with primers RE6490/RE6491.

The SNAP-eGFP fusion construct was created by amplifying the SNAP fragment from pIRES-PEX5-SNAP/eGFP-SKL with the primers KR011/KR012 and subcloning it into HindIII/BamHI digested peGFP-N1 (Clontech).

The PEX5L-SNAP fusion construct was created by amplifying the SNAP fragment from pIRES-PEX5-SNAP/eGFP-SKL with the primers KR022/KR026 and subcloning it into SalI/NotI digested peGFP-N1 (Clontech).

For the heterologous expression of eGFP fusion proteins in *E. coli,* the pET-9d (Merk, Darmstadt, Germany) vector was used. Here the N-terminal sequences of human PEX5L (amino acids 1-335) and PEX5 from *Trypanosoma brucei* (AA 1-340) were fused to eGFP using the FastCloning approach. The pET-9d His *Hs*PEX5(1-335) was amplified using the primers RE7008/RE7009 and eGFP was amplified with primers RE7010/ RE7011. This vector was then used to construct the pET-9d His *Tb*PEX5(1-340) eGFP by amplifying the backbone with the primers RE7012/RE7013 and the *Tb*PEX5(1-340) using the primers RE7016/RE7017.

### Recombinant proteins

For measuring the diffusion coefficients of the N-terminal halves of *Hs*PEX5L (AA1-335) and *Tb*PEX5 (AA1-340), both fused to eGFP, these were heterologous expressed in *E.coli* and purified using Ni-NTA columns as described elsewhere (Schliebs et al., 1999). Briefly, the cells were homogenized by sonication, sedimented and the supernatant was incubated with the Ni-NTA matrix for one hour. Afterwards, the proteins were eluted with a 10-500 mM imidazole gradient. For confocal FCS measurements, concentrations of 5 nM for *Hs*PEX5L N-Term and 20 nM for *Tb*PEX5 N-Term were used. eGFP was obtained from Novus Biologicals (Littleton, Colorado, US) and used at a final concentration of 10 nM.

### Subcellular fractionation to purify cytosol from human cells

Human HEK-293 cells were harvested from two T75 flasks for each experiment. The cells were incubated in 1 ml Digitonin buffer (150 mM NaCl, 50 mM HEPES, 25 μg/ml Digitonin) for 10 min, as described in (Holden and Horton, 2009). After sedimentation of the remaining cell fragments (2000 xg, 4 °C, 5 min), the cytosol was in the supernatant. For measuring the diffusion speed, the recombinant proteins were diluted with purified cytosol to 5 nM for His-*Hs*PEX5L (1-335) and 20 nM for His-*Tb*PEX5(1-340) and analyzed by point FCS.

### Construction of HEK-293 PEX5 and PEX14 knock out cell lines using CRISPR/Cas9

Deletion of PEX5 and PEX14 in HEK-293 cells was accomplished using a dual sgRNA approach. In particular, the sgRNA pairs targeted defined critical exons of PEX5 (exon 2) and PEX14 (exon 3), resulting in nonsense-mediated mRNA decay (NMD), the likely event after homozygous deletion in both cases. sgRNAs were selected using the CRISPOR algorithm (Haeussler et al., 2016). sgRNAs targeting the intron upstream of the respective splice branch point were cloned into pX458 (Addgene 48138, Dr. Feng Zhang), and sgRNAs targeting the intron downstream of the target exon were cloned into pX458-Ruby (Addgene 110164, Dr. Philip Hublitz). sgRNA ON-target efficiency was evaluated by Surveyor assay (Surveyor Mutation Detection Kit, IDT) and the following guides were chosen: PEX14 5’: GGatcagctcgaatggagatc, PEX14 3’: GGaccccccagtggggcatgc; PEX5 5’: ggggtcgcagcaaaagcact, and PEX5 3’: Ggtttataaacgctcagtaag (capitalized nucleotides within the sgRNA sequences correspond to added G residues to allow for proper pol III transcription). Cells were transfected with both sgRNA expressing plasmids. 72h post transfection double mRuby2/eGFP-positive cells were FACS-sorted into 96-well plates. After expansion, cells were analysed by genomic PCR using primers spanning the deleted region (PEX14 FW cagcatacagggcacaagggcgg, PEX14 RV: tgctactgaatgctgcctttgcc; PEX5 FW: ggtccaggcccctttgtggaggc, PEX5 RV: aacaagcaggcattctcattcgg). Mutations were verified by Sanger-sequencing of the subcloned amplicons, and absence of the respective protein was confirmed by immunoblotting. Of note, we obtained several clones in which the excised exon was found inverted and re-inserted, nevertheless giving a full KO phenotype as described (Blayney et al., 2020). All clones selected for this study were confirmed homozygous deletions.

### Sample preparation for live cell FCS measurements

Human fibroblasts (GM5756-T (RRID:CVCL_VQ75)) (South and Gould, 1999), HEK 293 (ATCC, VA, USA) HEK KO PEX5 and HEK KO PEX14 cells were maintained in a culture medium consisting of DMEM with 4500 mg glucose/L, 110 mg sodium pyruvate/L supplemented with 10% fetal bovine serum, glutamine (2 mM) and penicillin-streptomycin (1%). The cells were cultured at 37 °C/ 5% CO_2_. Cells were grown on #1.5 μ-Dish 35 mm (World Precision Instruments Sarasota, Florida, US) and transfected with 2.5 μg DNA per dish using Lipofectamine 3000 transfection reagent (Invitrogene, Carlsbad, USA). 24 hours after transfection, the cells were incubated at 37 °C for 40 min with SiRo snap-tag (1.2μM), SNAP-Cell® 647-SiR, or SNAP-Cell® 505-Star (both from New England Biolabs, Ipswich, MA, USA) to label the SNAP-tagged PEX5L. The samples were washed twice for 20 min with 1 ml of culture media in 37°C incubator. For the measurement the culture media was substituted with L-15 Medium (Leibovitz’s) (Thermo Fisher Scientific) media and placed on the microscope for data acquisition for no longer than 1 hour. Each sample was kept in culture media in the incubator until the measurement started.

### Confocal FCS measurements

#### Confocal live cell FCS measurements

Confocal FCS measurements on living cells were performed at a confocal Zeiss LSM 880 microscope equipped for fluorescence correlation spectroscopy. The microscope provides 488 nm and 633 nm excitation lines focused into the sample via a 40x/1.2 (Zeiss C-APOCHROMAT) water immersion objective lens. Excitations powers were set respectively for calibration solution and cytosolic measurements at 7 μW and 0.5 μW for the 488 nm excitation laser and at 6 μW and 1 μW for the 633 nm excitation laser (power measured at the objective lens). The fluorescence emission was split into two detectors depending on the characteristic emission wavelengths and the counts per molecule maximized acting on the correction collar of the objective lens. A time trace was recorded for each spectral emission channel and both fluorescence auto- and cross-correlation curves calculated using the Zen software (Zeiss).

In order to calibrate the illumination volume for both excitation lines, an 18 nM solution containing a 1:1 mix of Alexa Fluor 647 (330 μm^2^/s) (Eggeling et al., 2009b) and Alexa Fluor 488 (430 μm^2^/s) (Clausen et al., 2015) was used in a μ-slide 8 well dish (ibidi GmbH, Martinsried, Germany). The excitation beams were focused a few micrometres away from the glass of the microscope slides into the solution and an FCS measurement was repeated 3 times with a duration of 5 sec per time trace recording. The characteristic confocal full width at half maximum (FWHM), d, was calculated, knowing the diffusion coefficient, D, and deducing an average transit time, T_D_ from the 3 acquired FCS curves, related by Eq.1.

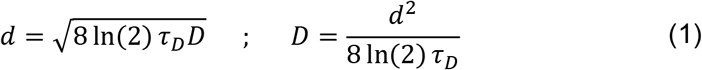

The calibrated d per excitation line was then utilized to calculate D for each protein of interest labelled with a dye with similar spectral characteristics, implying same excitation volume (eGFP and SiRo in our experiments). Each illumination volume was calibrated every day to monitor microscope performance and take small volume variations into consideration. Once the focal volume was calibrated the actual measurements on live cells could be performed. Healthy cells expressing eGFP and/or SNAP/Halo-tagged SiRo signal were selected via visual inspection of the samples. In each sample, 3 data collection locations in the cytosol of each cell were selected and the time trace acquisition repeated 3 times per each set position for a 5 sec recording. At least 10 cells for each sample were measured for the acquisition of one data set. No appreciable photobleaching occurred during the acquisition.

The Zeiss ZEN software provides already auto-correlated or cross-correlated curves that were analysed via FoCuS-point software (Waithe et al., 2016). The data were fit using a 3D diffusion model that includes a triplet component. The overall generic model for analysing the correlation curves was:

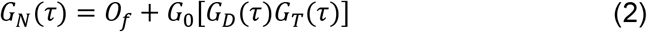

where t represents the correlation lag time, *O_f_* represents the offset (fixed to 1), *G_0_* is the amplitude of the correlation function at *τ* = 0, *G_D_(τ)* is describing all correlation components relating to diffusion processes and *G_T_(τ)* is an optional term accounting for a triplet state (dark state kinetics).

To analyse our data, we used the following triplet equation:

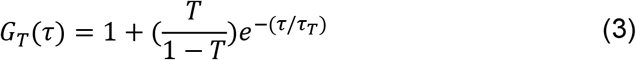

where T is the average triplet amplitude and *τ_T_* is the triplet correlation time that we fixed at 0.005 ms for SiRo and 0.04 ms for eGFP.

To analyse our cytosolic proteins’ mobility (G_D_(*τ*)) we considered a 3D diffusion model with multiple component fitting possibility:

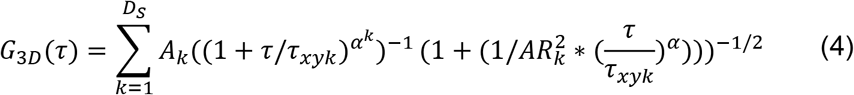

with alpha fixed to 1, the aspect ratio that describes the illumination volume AR=6 and *τ_xyk_* as the average transit time for the kth component with its fraction *A_k_*.

The diffusion analysis was optimized by comparing the diffusion of PEX5L and PEX5 S600W, both expressed in combination with eGFP-PTS1 in human cells. Here, the dual plasmids PEX5L-SNAP/eGFP-PTS1 and PEX5L S600W-SNAP/eGFP-PTS1 were expressed respectively. When PEX5L S600W-SiRo is expressed in human fibroblasts, the collected correlation curves for this protein were fitted considering one diffusing component, as the S600W substitution inhibits PTS1 binding and therefore the PEX5 S600W cannot bind/co-diffuse with the eGFP-PTS1. Only a free component of its characteristic transit time and the corresponding diffusion coefficient have been considered and calculated.

To study the dynamics of PEX5L, in cells expressing PEX5L-SNAP-SiRo, we initially introduced a two-component fitting model (D_S_=2 in the above model), trying to isolate a free diffusion contribution (that we could fix according to the previous analysis on PEX5L S600W) from a bound diffusion contribution (which we expected to diffuse together with PTS1 proteins). Clearly only one particular diffusion coefficient could be extracted from these data, comparable for PEX5L and PEX5 S600W, so the number of components that contribute to the fitting model related to PEX5L has been reduced to one. This also indicated that the binding of eGFP-PTS1 to PEX5L-SNAP-SiRo did not influence the diffusion speed of the receptor.

When PEX5 S600W was expressed in combination with eGFP-PTS1, the collected curves for the cargo protein eGFP-PTS1 were fitted considering two distinct populations (eGFP-PTS1 can still bind to the endogenous PEX5, fully functional but not labelled). The bound component was fixed in our analysis based on the PEX5L characteristic diffusion coefficient (recalculated accordingly for 488 nm illumination volume) and the PTS1 free diffusion component extracted from the acquired FCS curves as well as the proportion between bound and unbound fraction. The majority (more than 87% on average) of the PTS1 in these samples was found unbound.

In the case of eGFP-PTS1 expressed in combination with PEX5L (dual plasmid PEX5L-SNAP/eGFP-SKL), the FCCS curves showed a clear co-diffusion of PEX5L and eGFP-PTS1 at a diffusion coefficient comparable to that of PEX5L. Therefore, we considered the calculated PEX5L diffusion coefficient as the bound contribution. We set this diffusion coefficient as the characteristic diffusion coefficient of eGFP-PTS1 when bound to PEX5L (recalculated accordingly for 488 nm illumination volume). Finally, we combined the extracted information obtained by eGFP-PTS1 bound diffusion and free diffusion coefficients (from PEX5 S600W/eGFP-SKL expression) to calculate the fraction of eGFP-PTS1 diffusing free and bound in the case of the simultaneous expression of PEX5L-SNAP/eGFP-SKL.

Each calculated correlation curve was inspected by eye and eventually discarded when showing features clearly out of the expected fitting model. In most cases discarded measurements showed a bright spike in the time trace that biased the correlation curve, possibly caused by a fluorescent cluster or a whole peroxisome moving into the focal volume. We did not explore the PEX5L diffusion at the peroxisomal membrane, as there the protein mobility is slower than the photobleaching rate at the set conditions, which made it impossible to measure characteristic diffusion coefficients in this location.

### Proteins in solution

FCS measurements were performed at a confocal PicoQuant MicroTime 200 machine and a Zeiss 880 laser scanning microscope. The microscopes provided 488 nm excitation lines focused into the sample via a 60×/1.2 (Olympus UPlanSApo) and a 40×/1.2 (Zeiss C-APOCHROMAT) water immersion objective lenses, respectively. Excitation power was set to 5 μW to minimise eGFP photobleaching. Counts per molecule were maximized acting on the correction collar of the objective lenses. Time traces were recorded and fluorescence correlation spectroscopy curves calculated for each measurement.

To measure the diffusion of eGFP and N-terminal part of PEX5L (human and from *Trypanosoma brucei,* fused to eGFP) in solution, the proteins were diluted to ca. 10 nM. For each condition, recordings at 3 different spots were performed by measuring for 15-20 s 5 times at about 10 μm depth from the coverslip. Time traces acquired at the MicroTime were correlated using the FoCuS-point software. Zeiss data were correlated using the ZEN software. The data were fitted using a 3D diffusion equation that includes a triplet component as described in the previous section.

### AO-zSTED-FCS measurements

We used a custom STED microscope built around a RESOLFT microscope from Abberior Instruments as described previously (Barbotin et al., 2019b; Galiani et al., 2016). The microscope was equipped with a 640 nm pulsed excitation laser focused into the sample via a 100x/1.4 (Olympus UPLSAPO) oil immersion objective lens. Excitation power was set to 6 μW (measured at the back aperture of the objective lens) in cells. A 755 nm depletion beam pulsed at 80 MHz (Spectra-Physics MaiTai, pulse-stretched by a 40 cm glass rod and a 100 m single-mode fibre) was modulated in phase using a spatial light modulator (SLM) (Hamamatsu LCOSX10468-02). STED power was set to 16 mW for the aberration correction procedure and varied between 6 and 33 mW for STED-FCS measurements. The fluorescence emitted by the sample was collected back by the objective lens, filtered via emission filters and a pinhole and detected using an avalanche photodiode (APD). The system was equipped with a correlation card (Flex02-08D) to acquire both time traces and FCS curves. The microscope was controlled by the Imspector software (Abberior Instruments). The SLM was employed for both phasemask generation and aberration correction and controlled by a bespoke python software as described in (Barbotin et al., 2019a). z-STED illumination volume calibration was carried out on a bead sample, FluoSpheres, crimson (625/645), diameter = 0.04 μm.

On each studied sample an aberration correction procedure was run at the beginning of any data acquisition on one selected cell at approximately 3 μm from the slide into the cytosol. Data collection consisted of a series of time traces acquired at increasing STED power distributed over a z-STED beam. For each sample at least 5 cells were selected and a STED power series (0, 7, 16, 33 mW) was run over a selected point in the cytosol. Each measurement was acquired for 10 sec and repeated 3 times. The data was fitted using a standard 3D diffusion equation reported above (Eq 4), with an alpha parameter set to 0.75 to account for diffusion in the crowded environment of the cytoplasm (Barbotin et al., 2019b; Weiss et al., 2004). To account for the simultaneous decrease in lateral and axial size of the observation volume, we fitted STED-FCS curves using the model we previously developed (Barbotin et al., 2019b). In short, confocal FCS curves were fitted first with an aspect ratio set to 4 (different from above as a higher-NA objective was used here). For z-STED recordings, the relative decreases in aspect ratios and lateral transit times with respect to the confocal values were fitted together using a single parameter, from which the axial transit times (t_z_) were calculated. The relationship between lateral and axial dimension of the observation volume was calibrated using images of fluorescent beads, as described in (Barbotin et al., 2019a). No appreciable photobleaching occurred during the acquisition.

## Acknowledgements

We thank the Wolfson Imaging Centre Oxford for providing microscope facility support for data acquisition and analysis as well as the Microverse Imaging Center Jena. We acknowledge funding by the Wolfson Foundation, MRC (Grant No. MC_UU_12010/unit programs G0902418 and MC_UU_12025), the Wellcome Trust (grant No. 104924/14/Z/14, Strategic Award 091911 (Micron)), MRC/BBSRC/EPSRC (Grant No. MR/K01577X/1, MRC grant number MC_UU_12010/unit programmes G0902418 and MC_UU_12025), the EPA Cephalosporin Fund, the John Fell Fund, and the Deutsche Forschungsgemeinschaft (Research unit 1905 “Structure and function of the peroxisomal translocon”; grant number 322325142 “Super-resolution optical microscopy studies of peroxisomal protein import in the yeast Saccharomyces cerevisiae”, Germany’s Excellence Strategy – EXC 2051 – Project-ID 390713860, project number 316213987 – SFB 1278). P.C. acknowledges a postdoctoral fellowship from the Basque Government (POS_2018_1_0066 and POS_2019_2_0022).

## Author contributions

SG, KR, AB, PC, IU, JO, JK, PH performed the experiments. ES, FS, DW helped analysing the FCS data. WS, RE, CE gave input on the experimental design.

## Abbreviations list

ACF: autocorrelation function
AO: Adaptive Optics
CRISPR: Clustered Regularly Interspaced Short Palindromic Repeats
FCS: Fluorescence Correlation Spectroscopy
GPMV: Giant Plasma Membrane Vesicle
PEX: peroxin
PTS1: peroxisomal targeting signal type 1
SiRo: Silicon Rhodamine
STED: stimulated emission depletion

